# Low-Density Lipoprotein Receptor-Mediated Lipidome-Transcriptome Reprogramming Impulses to Cisplatin Insensitivity

**DOI:** 10.1101/502401

**Authors:** Wei-Chun Chang, Hsiao-Ching Wang, Wei-Chung Cheng, Juan-Cheng Yang, Wei-Min Chung, Lumin Chen, Yen-Pin Ho, Yao-Ching Hung, Wen-Lung Ma

## Abstract

Platinum-based therapy remains the cornerstone for cancer patient management; however, its efficacy varies. Theis study demonstrated the differential expressions of low-density lipoprotein receptor (LDLR) in subtypes of epithelial ovarian carcinoma (EOC) determines cisplatin sensitivity. It’s sensitive in serous EOCs (low LDLR), where insensitive in endometrioid and clear cell EOCs (high LDLR). Meanwhile, knocked-down or overexpressed LDLR in EOC could reversed the chemosensitivity pattern both in vitro and in vivo. Mechanistic dissection with transcriptome vs. lipidome trans-omics analyses elucidated the LDLR→LPC (Lyso-PhosphotidylCholine)→FAM83B (phospholipase-related)→FGFRs (cisplatin sensitivity and phospholipase-related) regulatory axis in cisplatin insensitivity. Implementing LPC-liposome encapsulated cisplatin could facilitate DNA-adduct formation via lipid droplets (LDs) delivery. Furthermore, Bioinformatics analyses found that the LDL/R→LD homeostasis alteration is critical for therapeutic prognosis. Lastly, using LPC-liposome-cisplatin improved cisplatin sensitivities in gastric cancer, renal cell carcinoma, hepatocellular carcinoma, cholangiocarcinoma, and pancreatic adenocarcinoma cells. In conclusion, this report discovered a LDL/R-reprogrammed transcriptome-lipidome network, by which impulses platinum insensitivity and disease outcome. The drug specific lipidome for liposome manufacture might be an efficienct pharmaceutics strategy for chemoagents.

**Significance:** LDLR reprograms cellular lipidome and transcriptome profiles to determines chemotherapy therapeutic efficacy. The LDLR-reduced LPC abundance disturbs phospholipids homeostasis of Lands cycle in LD, by which attenuates intracellular platinum transportation for DNA-adduct formation. Targeting LDLR-LD-lipidome with LPC-liposome-platinum could boost therapeutic efficacy for insensitivity.

## Introduction

Platinum-based adjuvant chemotherapy is widely used in the management of patients with solid tumors, including gynecological malignancies (Falcetta, Medeiros et al., 2016, Lawrie, Winter-Roach et al., 2015), gut cancers (Galdy, Cella et al., 2016, Ishikawa, Abe et al., 2016, Malka, Cervera et al., 2014), urinary tract carcinoma (Roupret, Neuzillet et al., 2016), and non-small cell lung cancer (Berghmans, Scherpereel et al., 2017, de Castria, da Silva et al., 2013) and many other malignancies. Platinum can form DNA adducts in fast growing cancer cells, a capacity which makes it an excellent tumor growth suppressing agent. However, while platinum-based chemotherapies are commonly used to treat human malignancies, variations in the responsiveness and resistance of patients to such therapies are also seen (Bellmunt, Pons et al., 2013, Johnson, Bryant et al., 2011, Poonawalla, Parikh et al., 2015). The mechanisms underlying platinum-based chemotherapy sensitivity and resistance have yet to be elucidated; however, there is an interesting correlation between histological cellularity and platinum therapy responsiveness (Falcetta et al., 2016, Ferreira, Peixoto et al., 2016, Glasspool & McNeish, 2013, Prendergast, Holzapfel et al., 2017).

Among various solid tumors, epithelial ovarian carcinomas (EOCs) arise from diversified origins of epithelium including ovarian epithelial cells, the fallopian tubes, and cells that have migrated from endometriosis or the intestines (Klotz & Wimberger, 2017, Vargas, 2014). Despite inconclusive evidence regarding the origins of EOCs, however, dissecting the complexity of EOC cellularity with chemotherapy sensitivity might clarify why variations in platinum responsiveness among EOCs occur (McCluggage, 2011). The different subtypes of EOCs can be classified histologically as serous, mucinous, endometrioid, and clear-cell EOCs (Vargas, 2014), and previous studies have found that these subtypes of EOCs are associated with varying levels of platinum therapy efficacy (Dahm-Kahler, Borgfeldt et al., 2017, Glasspool & McNeish, 2013, Lawrie et al., 2015, Ledermann, Luvero et al., 2014, Matulonis, Sood et al., 2016). The serous EOC cells/patients have good sensitivity to chemotherapy (McCluggage, 2011). In contrast, advanced clear-cell EOCs have been found to exhibit poor responses to platinum-based chemotherapy, with recurrent clear-cell carcinomas appearing to be particularly resistant to chemotherapy and difficult to treat (Lee, Kim et al., 2011, Mizuno, Kikkawa et al., 2006, Sugiyama, Kamura et al., 2000). Malignant mucinous tumors are epithelial ovarian tumors formed by cells that resemble the endocervical epithelium or intestinal epithelium (intestinal type), and the efficacy of platinum-based therapies against these tumors is unknown (Brown & Frumovitz, 2014). Meanwhile, the clinicopathological features of malignant endometrioid EOCs are similar to those of clear-cell EOCs, as these subtypes are postulated to arise from the same cell type (Shevchuk, Winkler-Monsanto et al., 1981) and to be platinum-based chemotherapy insensitive (Sugiyama et al., 2000).

Lipids are essential for biomass and building block synthesis, and lipids can also act as bioactive molecules, e.g., as constituents of cellular membranes, or as an energy supply sufficient for the fast-growing nature of cancer cells (Ward & Thompson, 2012). Microscopically, endometrioid EOC exhibits an appearance similar to that of tubular glands and bears a resemblance to endometrium. Squamous differentiation is commonly seen in endometrioid EOC patients (Wagner, Buck et al., 1994), along with surrounding lipid droplet (LD)-vacuolated stromal tissue (Ulbright & Roth, 1985). In addition, clear-cell EOC is also characterized by a significant amount of LD vacuoles in the cytoplasm (Nishimura, Tsuda et al., 2010). Moreover, the lipophilic nature of metastatic EOCs cells keen to migrating to the omentum (Nieman, Kenny et al., 2011). All of those characteristics have raised the question of whether non-autonomous lipid providers, i.e., tumor macroenvironmental regulators (Lee, Chang et al., 2016) produced via the lipoprotein-mediated lipid route (lipoprotein/receptor-route) (Chang, Huang et al., 2017), might play roles in the development and disease progression of EOCs.

In the current study, we discovered that different levels of low-density lipoprotein receptor (LDLR; the gate for non-autonomous lipid entrance) expression in EOCs determine platinum sensitivity in an LDLR-dependent manner. In addition, LDLR alters the lipid profile and LD homeostasis of EOC platinum therapy sensitivity. Lastly, the targeting of the LDLR-lipidome and its effects on cisplatin sensitivity can be generalized in multiple cancer types.

## Results

### LDLR expression is platinum chemosensitivity confounder

Given our hypothesis that the entry of tumor macroenvironmental lipid through the lipoprotein/receptor-route might contribute to the development of EOC subtypes, we examined the expression of LDL/R-route related genes (Chang et al., 2017) in EOC patients. The LDL/R-route components include LDLR (the entrance gate for LDL) and LPL (lipoprotein lipase; which is responsible for unloading lipid from LDL). As indicated by the results shown in Fig. 1A, 1B, and 1C, LDLR and LPL were expressed throughout the various EOC subtypes. However, predominantly weak LDLR staining was seen in serous and mucinous EOC patients, whereas predominantly strong LDLR staining was seen in endometrioid and clear-cell EOC patients. Compared to that for LDLR, the expression pattern of LPL was more consistent across the various subtypes of EOC. Examining the LDLR expressions in EOC cells originated from serous (OVCAR3, SKOV3), endometrioid (MDAH-2774, TOV-112D), and clear-cell (TOV-21G, ES2) EOCs, we found that the endometrioid and clear-cell EOCs had abundant LDLR (Fig. 1D). The serous EOC was the subtype most sensitive to cisplatin, with a low IC 50 value compared to those for the endometrioid and clear-cell EOCs (Fig. 1E). A similar phenomena was observed in the colony formation assay results (Fig. 1F). Overall, the results shown in Figure 1 demonstrated high degrees of correlation among the subtypes of EOCs, their LDLR expressions, and their cisplatin sensitivities.

**Figure 1.**
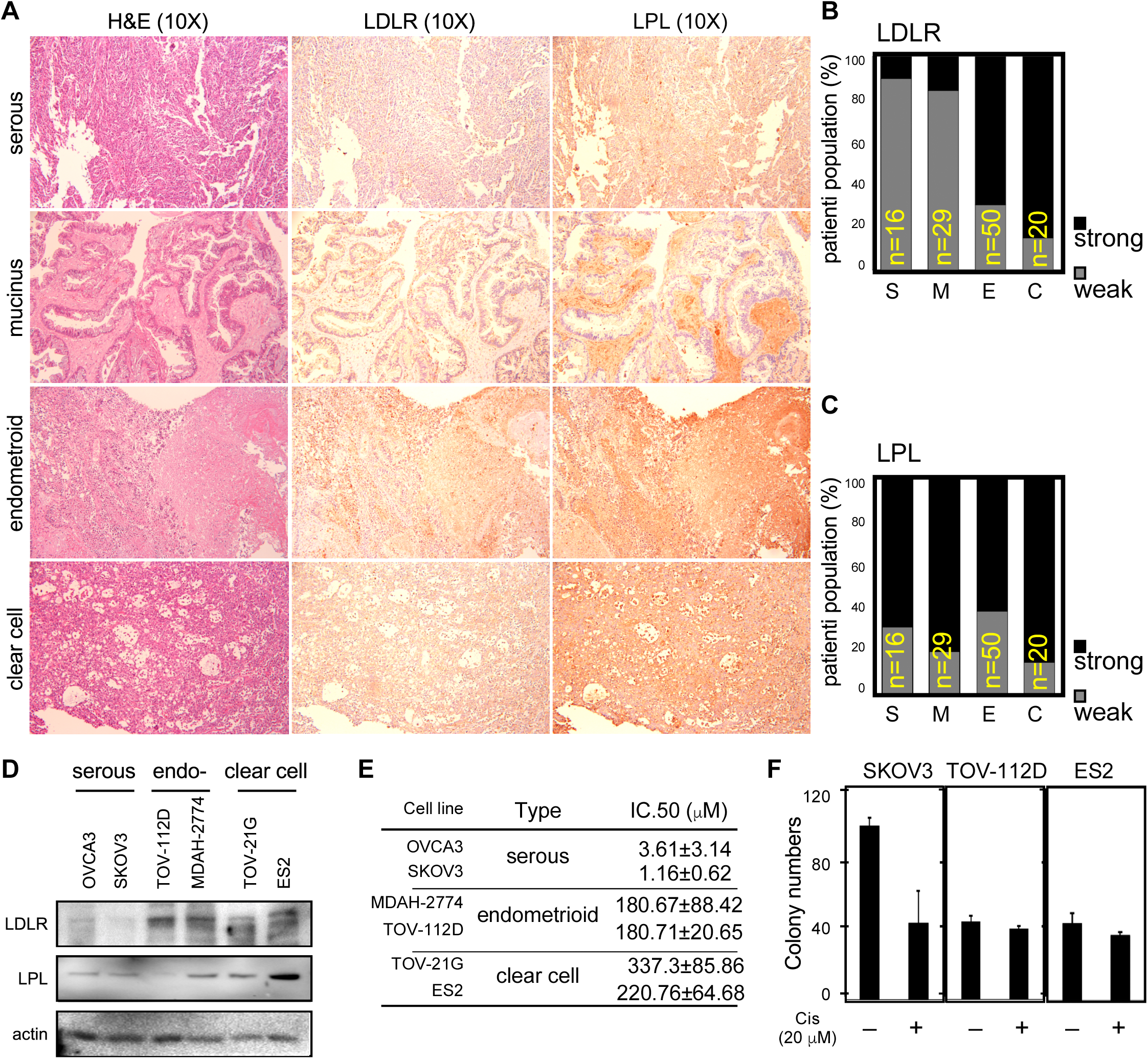
Differential expressions of LDLR in EOC subtypes. **A**. Histological H&E staining (left column) and immunohistochemistry (IHC) staining of LDLR (middle column) and LPL (right column) of serous EOC (1^st^ row), mucinous EOC (2^nd^ row), endometrioid EOC (3^rd^ row), and clear cell EOC (4^th^ row) cells. **B**, **C**. Quantitation of IHC scores of LDLR (B) and LPL (C) in EOC subtypes. The patient numbers are serous (S; n=16), mucinous (M; n=29), endometrioid (E; n=50), and clear cell (C; n=20). “Strong” indicates an IHC score of 3 or above, while “weak” indicates an IHC score lower than 3. **D**. Differential LDLR (upper panel) and LPL (middle panel) in EOC cells. S: OVCAR3, SKOV3; E: MDAH-2774, TOV-112D; C: TOV-21G, ES2. Actin served as the loading control (lower oanel). **E**. Cisplatin cytotoxic IC. 50 value of EOC cells. The unit of the IC. 50 values is μM. **F**. Differential inhibitory efficacies of long-term (14 days; colony forming capacity), low-dose (20-μM) cisplatin treatments on SKOV3, TOV-112D, or ES2 cells. Upper panels are representative images of colonies on plates, and lower panels show the quantitation of colony assays. All in vitro data are from at least three representative experiments.

In order to test the role of LDLR expression in cisplatin responsiveness *in vitro*, shRNA targeting LDLR expression was introduced in endometrioid and clear cell EOCs, while stably transfected LDLR cDNA was introduced in serous EOC (Fig. 2A). We found the LDLR cDNA could reduce while the shRNA could enhance cisplatin colony suppressing capacity (Fig. 2B) and cytotoxic efficacy (Fig. 2C). The IC 50 value of cisplatin could be significantly altered in TOV-112D (from 427±27.4 to 111±32.6 μM), TOV-21G (from 178±72.7 to 99±33.7 μM), MDAH-2774 (from 125±26.6 to 63±0.5 μM), and SKOV3 (from 8.9±1.06 to 297.2±15.43 μM) cells (Fig. 2D). To further test the effects of LDLR expression on cisplatin responsiveness *in vivo*, we performed the xenograft tumor with cisplatin treatment procedure (see the methods section) in mice bearing MDAH-2774, TOV-21G, and SKOV3 EOC cells. We found that LDLR knockdown in the tumor with shLuc-infected MDAH-2774 cells could suppress the effects of cisplatin treatment to a minor degree; however, tumors with LDLR knockdown tumors caused by shLDLR were dramatically ameliorated by the same treatment procedure (Fig. 3A). In contrast, the tumor with pLenti-infected SKOV3 cells could be inhibited by the cisplatin treatment procedure, and the effects of treatment were comparable in tumors in which LDLR cDNA was stably expressed (Fig. 3B). To further confirm the effects of LDLR expression on cisplatin responsiveness, we performed the same procedure in tumors with shLuc-infected TOV-21G cells and found very little response. However, the tumors with shLDLR-infected TOV-21G cells could be almost abolished with the treatment procedure. To sum up the results shown in Fig. 1, Fig. 2, and Fig. 3, the expression levels of LDLR are the key cisplatin insensitivity confounder.

**Figure 2.**
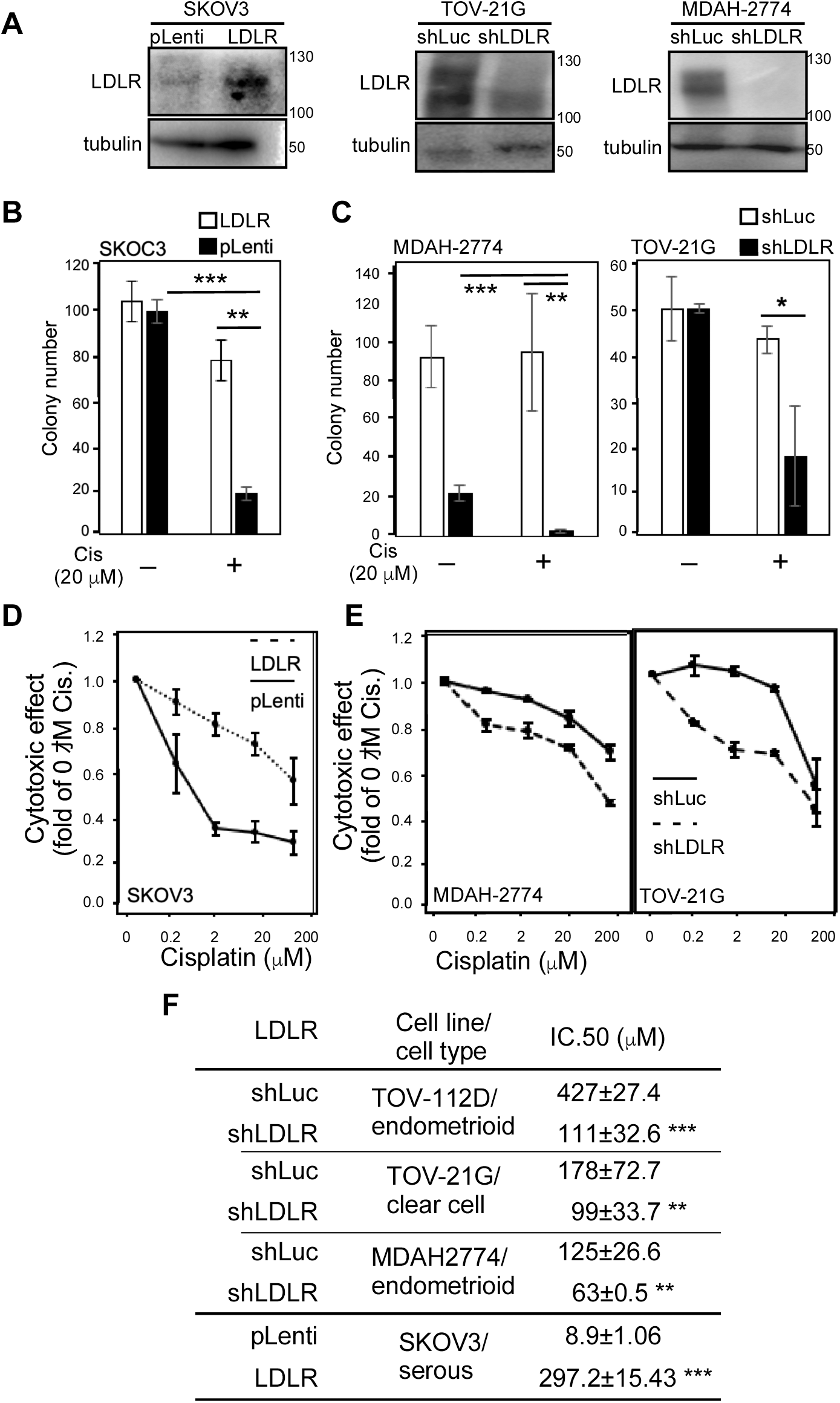
The expressions of LDLR determine cisplatin sensitivity in EOC cells. **A**. Manipulation of LDLR expressions in EOC cells. Left-hand side panel: stable transfection of LDLR cDNA in SKOV3 cells. pLenti: control empty vector; LDLR: pLenti-LDLR cDNA. Middle panel: stable transfection of shRNA targeting LDLR in TOV-21G cells. shLuc: pLKO.1 vector constructed shRNA targeting luciferase gene;shLDLR: pLKO.1-shLDLR targeting LDLR expression. Right-hand side panel: stable transfection of shRNA targeting LDLR in MDAH-2774 cells. **B**. Cell growth inhibitory efficacy of long-term, low-dose cisplatin treatments (Wang et al., 2018) on parental and stable LDLR expression SKOV3 cells. **C**. Cell growth inhibitory efficacy of long-term, low-dose cisplatin treatments on shLuc and shLDLR knockdown TOV-21G (left-hand side panel) and MDAH-2774 (right-hand side panel) cells. **D**. Cytotoxic effect of cisplatin on parental and stable LDLR expression SKOV3 cells. **E**. Cytotoxic effect of cisplatin on shLuc and shLDLR knockdown TOV-21G (left-hand side panel) and MDAH-2774 (right-hand side panel) cells. **F**. LDLR expressions determine cisplatin cytotoxic IC 50 values in various subtypes of EOC cells. The unit of the IC 50 values is μM. The ** indicates a significant difference due to a p-value < 0.01, and *** indicates a p-value < 0.001.

**Figure 3.**
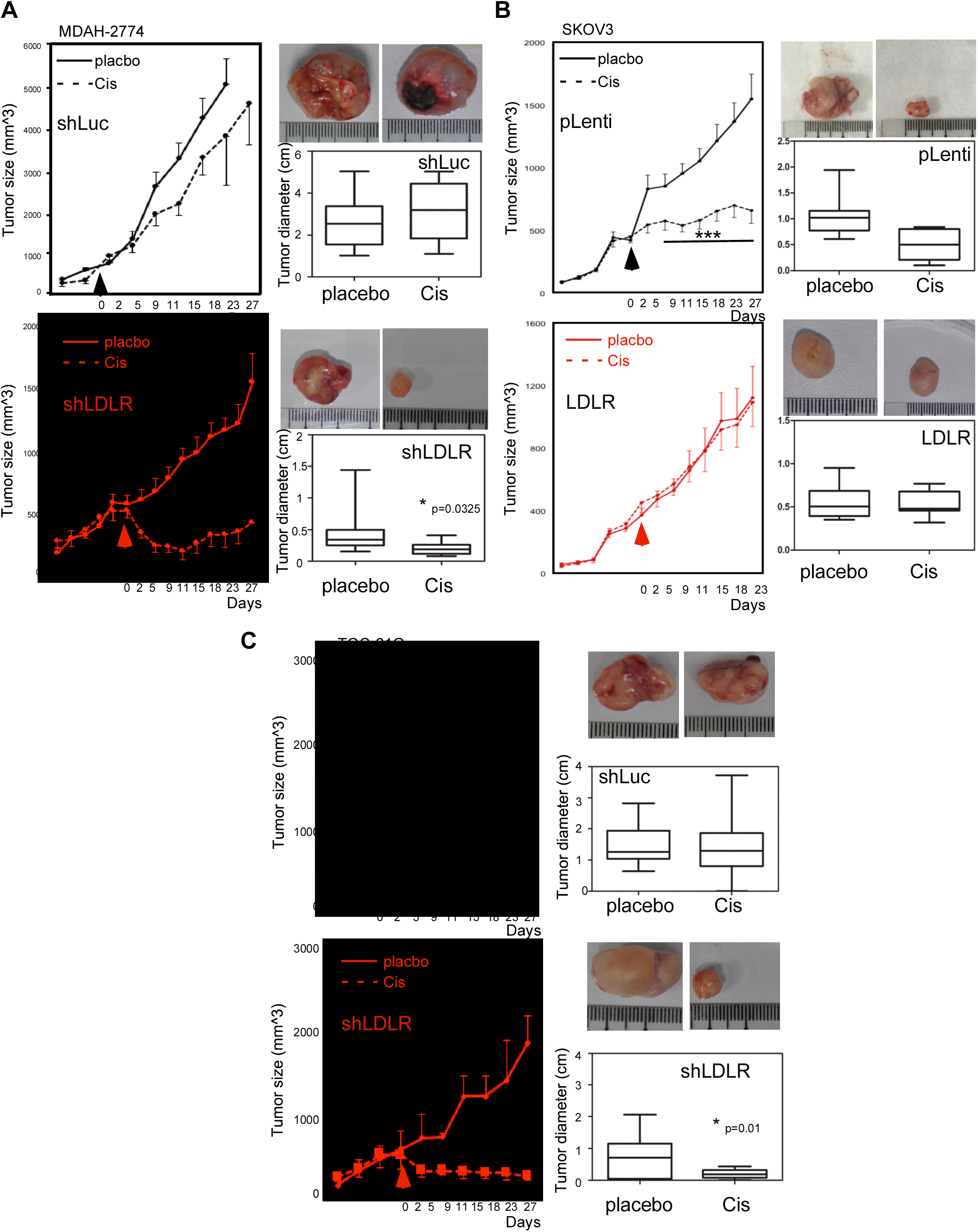
The expressions of LDLR determine cisplatin sensitivity in EOC xenograft tumors. Tumor suppressive effect of cisplatin (6 mg/kg/mice) in EOC cell (**A**. MDAH-2774; **B**. SKOV3; **C**. TOV-21G) xenograft mouse models. The left-hand side panels of each EOC tumor model show the tumor growth curve. The mice received cisplatin treatment (I.P.; 3 times/week for 3~4 wks) while the given tumor grew to 500 mm^3^. The right-hand side panels show a representative image (upper-right) and quantitation (lower-right) of the tumors. The cisplatin therapy efficacy levels of shLuc (upper panels) and shLDLR (lower panels) in mice bearing MDAH-2774 (**A**) and TOV-21G cells (**C**) were compared, as were those of pLenti- (upper panels) and LDLR (lower panels) in mice bearing SKOV3 cells (**B**). The * indicates a significant difference due to a p-value < 0.05, and *** indicates a p-value < 0.001. Each experimental group contained 10 mice with 20 tumor sites.

### LDLR→LPC→FAM83B→FGFR3 regulatory axis in platinum sensitivity

In order to determine the molecular mechanism of LDLR-mediated cisplatin insensitivity, we subjected two types of EOC cells (MDAH-2774 and TOV-21G; control vs. knockdown LDLR) to transcriptome profiling with RNA next-generation sequencing technology (RNAseq). As we aligned the transcriptome profile, we found that 1404 genes were consistently altered (Fig. 4A). Through gene ontology-based annotation and functional enrichment analysis, we were then able to determine the top 10 enriched pathways in terms of molecular function, which are listed in Fig. 4B. It was surprising to find that the lipid metabolism-related pathways were not prioritized in the LDLR knockdown list, whereas transmembrane receptor activity and transcription factor binding activity were. In verifying the LDLR knockdown effect on chemoresistant-related growth factor receptors, we found FGFR1~FGFR3 mRNA were significantly decreased by LDLR knockdown (Fig. 4C). Furthermore, the downstream signaling of FGFRs, including the phosphorylation of FAK and MEK, was also suppressed (Fig. 4D).

**Figure 4.**
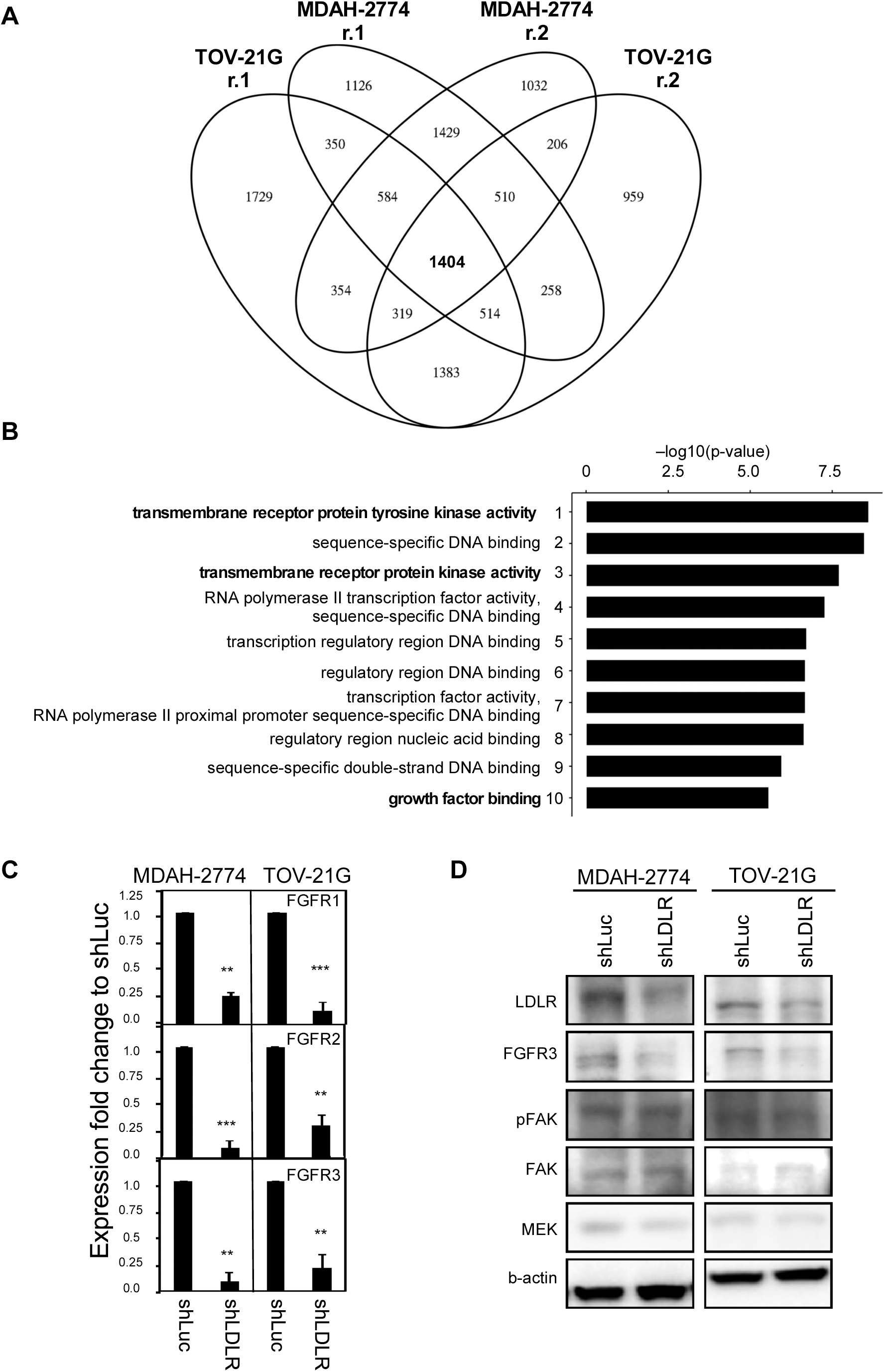
LDLR reprogrammed cellular transcriptome effects on receptor tyrosine kinase (RTK) and gene transcription activities. **A**. Replicated transcriptome analysis by RNAseq was performed to compare shLuc vs. shLDLR in MDAH-2774 and TOV-21G cells. The overlapped transcriptome showed that 1404 genes were consistently altered. **B**. Pathway enrichment analysis by GO-term followed with GSEA analysis. The top-10 enriched pathway were ranked from highest to lowest P-value(-log10). **C**. The mRNA expression of selected RTK (FGFR1~FGFR3) were measured in MDAH-2774 (left-hand side) and TOV-21G (right-hand side) cells comparing shLuc vs. shLDLR. **D**. The RTK downstream signaling, e.g., as indicated by pFAK/FAK and MEK amounts, was measured. The representative blots are shown, and beta-actin served as the loading control.

Considering the nature of LDLR for lipid importing, we performed lipidome analysis of three subtypes of EOC cells using shotgun lipid profiling technology. The lipid profiles were differentially expressed in six EOC cell lines, including with specific preferences (Fig. 5A). The ether-linked phospholipids were highly expressed in serous and endometrioid EOC cells. On the other hand, the storage lipids (DAG and TAG; which exist in endoplasmic reticulum and LD) were predominantly expressed in endometrioid and clear-cell EOC cells. To determine the LDLR knockdown effect on lipidomes, we performed lipid profiling (of MDAH-2774 and TOV-21G cells) comparing control vs. LDLR knockdown cells. The analyses revealed that lyso-phosphatidylcholine (LPC; up-regulated; EV1) and ether-linked phosphatidyletholamine (PE O-; down-regulated; EV 1) were significantly altered by LDLR knockdown (Fig. 5B; and Expanded Table 1).

**Figure 5.**
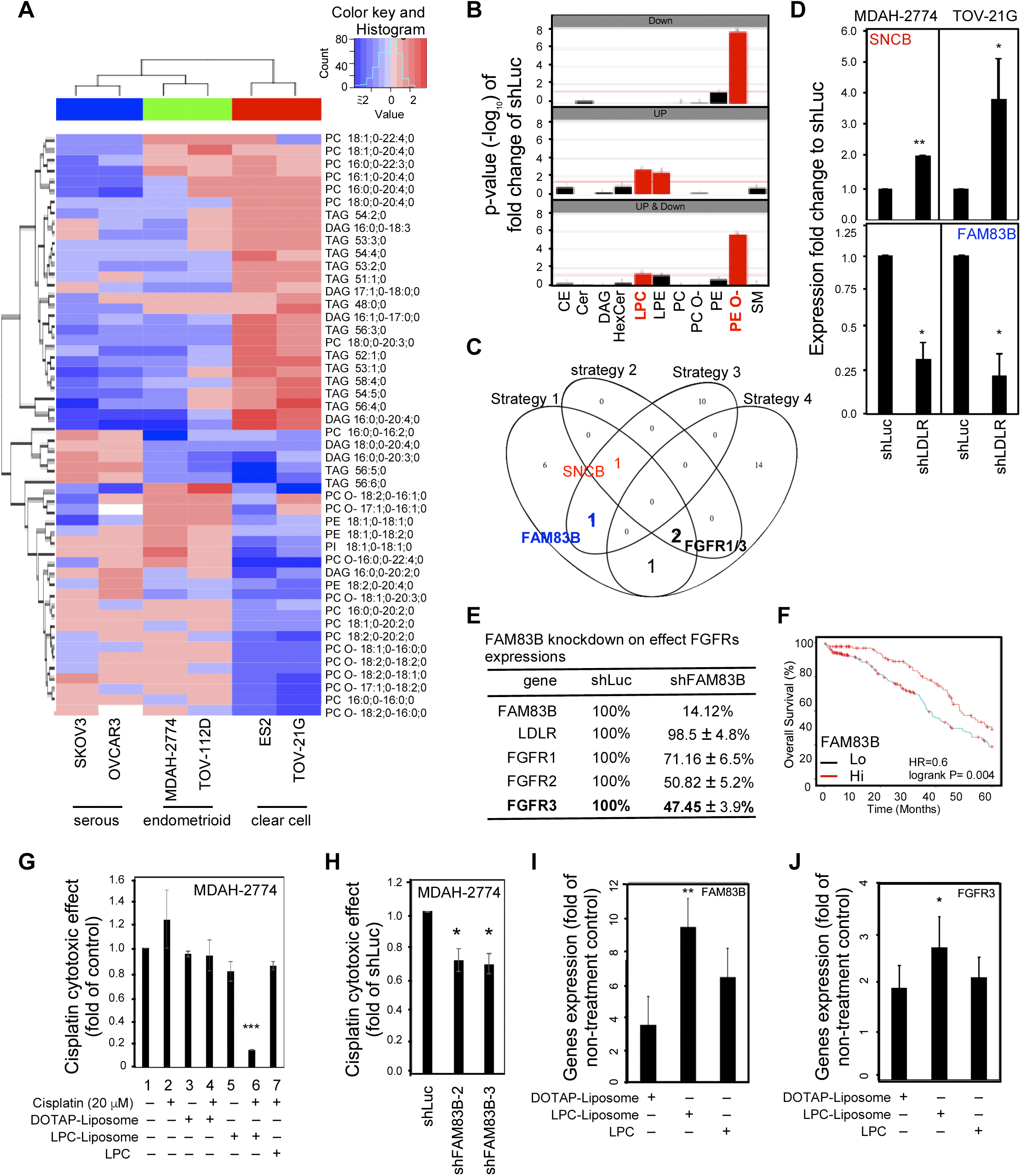
LDLR→LPC→FAM83B→FGFR3 regulatory axis in cisplatin insensitivity. **A**. Heat-map lipid profiling of six types of EOC cells, including serous (SKOV3, OVCAR3), endometrioid (MDAH-2774, TOV-112D), and clear cell (ES2, TOV-21G) lines. The spectrum from blue to red indicates the variation of lipid species among the cells. **B**. Lipidome analyses of lipid species comparing shLuc vs. shLDLR in MDAH-2774 and TOV-21G cells. The Y-axis shows the lipid species change p-value, where the threshold > −log_10_(1.4) indicates significant alteration (red-colored). LPC is increased and PE O- is decreased when the LDLR is knocked down. **C**. Trans-Omics analyses of transcriptome and lipidome profiles. The four selection criteria were implemented in the analyses. Strategy 1:. transmembrane receptor activity genes revealed by transcriptome analyses; Strategy 2: lyso-phospholipid-related phospholipase genes revealed in transcriptome analyses; Strategy 3: phospholipase expression negatively correlated with LDLR expression (for knockdown experimental design); Strategy 4: phospholipase genes significantly correlated to cancer survival revealed in TCGA database. The results came out SNCB (1 ∩ 2 ∩ 3), FAM83B (1 ∩ 3), FGFR1/3 (2 ∩ 4), and FGFR1 (1 ∩ 4). **D**. Confirmation of LDLR knockdown effect on SNCB (panels on left-hand side) and FAM83B (panels on right-hand side) mRNA expression. The SNCB and FAM83B mRNA expression were measured in MDAH-2774 and TOV-21G shLuc vs. shLDLR cells. **E**. Knockdown FAM83B effect on LDLR, FGFR1~3, etc. mRNA expressions. Knocking down FAM83B reduced the expressions of FGFR2 and FGFR3 and slightly decreased the expression of FGFR1, but did not alter LDLR mRNA expression. **F**. FAM83B is negatively correlated to EOC patient overall survival. The DriverDB.v2 platform was used to analyse TCGA data regarding the 5-year overall survival of EOC patients 5-years. The Hazard ratio was 0.6, and p = 0.004. **G**. LPC-liposome boosted cisplatin cytotoxic efficacy. Various treatments, e.g., w/ cisplatin vs. w/o cisplatin (lane 1 vs. 2), DOTAP-liposome (w/ cisplatin vs. w/o cisplatin 20 μM; lane 3 vs. 4), LPC-liposome (w/ cisplatin vs. w/o cisplatin 20 μM; lanes 5 vs. 6), and co-treatment with LPC and cisplatin (lane 7), were tested. **H**. FAM83B knockdown enhanced the cytotoxic effect of cisplatin in MDAH-2774 cells. The cytotoxic effect of cisplatin in shLuc cells was compared with two clones of shFAM83B (shFAM83B-2 and shFAM83B-3). **I**. The LPC-liposome upregulated FAM83B mRNA expressions. **J**. The LPC-liposome upregulated FGFR1~3 mRNA expressions. The * indicates a p-value less than 0.05; ** indicates a p-value less than 0.01; and *** indicates a p-value less than 0.001. The lipid profiling data were from two replicated experiments, and the gene expression and cytotoxic assay results were from at least three independent experiments.

Since lysophatidyl-lipids and phosphatidyl-lipids are the important lipids for LD metabolism (i.e., in the Lands cycle, in which they are catalyzed by LPCAT1/2/3 and PLA2) (Moessinger, Klizaite et al., 2014) and differ in different lyso-groups, we performed trans-omics analysis within transcriptome vs. lipidome data. There were four strategies applied in the analysis: 1^st^. transmembrane receptor activity genes revealed by transcriptome analyses; 2^nd^. lyso-phospholipid related phospholipase genes revealed in transcriptome analyses; 3^rd^. phospholipase expression negatively correlated with LDLR expression (for knockdown experimental design); 4^th^. phospholipase genes significantly correlated to cancer survival revealed in TCGA (The Cancer Genome Atlas) database.

The results shown in Fig. 5C indicated that there were no genes identified via all four strategies. Interestingly, however, there was one gene identified via strategies 1, 2, and 3 (SNCB, synuclein B; upregulated); one identified via strategies 1 and 3 (FAM83B, family with sequence similarity 83 member B; downregulated); and one identified via strategies 1 and 4 (VEGFR; downregulated). There were two genes identified via strategies 1, 2, and 4 (FGFR1 and FGFR3; upregulated), and their expression are reversed correlated to LDLR in patients (EV 1B; TCGA database). Those analysis results suggested an indirect regulation of phospholipase, LDLR, and RTK activity in EOC cells. In order to verify the correlations of regulation among gene expressions, we examined the LDLR knockdown effect on SNCB and FAM83B. The results were consistent with the transcriptome (Fig. 5D) in that SNCB was upregulated and FAM83B was downregulated by LDLR knockdown. Moreover, while the knockdown of FAM83B in MDAH-2774 cells mimicked the relationship between LDLR and FGFRs, the FGFR1~3 expression was significantly decreased by FAM83B knockdown (Fig. 5E). In brief, the mechanistic dissection revealed an LDLR→FAM83B→FGFR3 axis. Further examination using TCGA database with DBdriver.v2 analysis found a survival benefit of FAM83B expression in EOC patients in terms of 5-year overall survival (Fig. 5F), a finding which supports the LDLR→FAM83B→FGFR3→platinum therapy response axis observed *in vitro*.

We then tested whether the LDLR-mediated FAM83B→FGFRs regulatory axis affects platinum sensitivity in the context of an LPC-associated event. We compared the cytotoxic effects of direct treatment with cisplatin, treatment with liposome-encapsulated cisplatin (EV2; DOTOP-liposome; 1,2-dioleoyl-3-trimethylammoniumpropane), and treatment with LPC-liposome-encapsulated cisplatin on the insensitive MDAH-2774 cells. Only 20 μM of cisplatin encapsulated in LPC-liposome exhibited excellent cytotoxic efficacy (Fig. 5G; lane 1 vs. 5 vs 6). However, neither LPC-cisplatin co-treatment (non-liposome) (Fig 5G; lane 6 vs. 7; and EV3) nor DOTAP-liposome-cisplatin (Fig. 5G; lane 3 vs. 4) showed such efficacy. When the effect of FAM83B knockdown on cisplatin cytotoxicity was tested, it was found that two clones of FAM83B knockdown could significantly facilitate cisplatin cytotoxic activity in MDAH-2774 cells (Fig. 5H). In testing the LPC-liposome effect on gene expressions, meanwhile, we found that both FAM83B (Fig. 5I) and FGFRs (Fig. 5J) were upregulated in comparison with DOTOP-liposome. In brief, the results of the mechanistic dissection shown in Fig. 4 and Fig. 5 revealed an LDLR→LPC →FAM83B→FGFR3→platinum insensitivity axis.

This LDLR→LPC→FAM83B →FGFR3→platinum insensitivity axis had thus been delineated at the molecular level using trans-omics and in vitro analyses. However, the role it plays in patients remained unknown, as did its effects on the lipidome at the cellular and organelle levels. Using KMplotter survival analyzer, we found that high LDLR expression is associated with the poor prognosis of endometrioid EOC patients in terms of 5-year overall survival (Fig. 6A). We further found that LPL expression is associated with the poor prognosis of platinum-based (Fig. 6B) and taxol-based (Fig. 6C) treatments in EOC patients. These findings suggested that the LDL/R-route-chemotherapy sensitivity relationship is consistent with laboratory findings. In evaluating the LDLR→lipidome relationship in patients, and considering that the phosphatidyl-lipids vs. lyso-phosphatidyl-lipid conversion (Lands cycle) takes place in LDs, we hypothesized that the LDL/R-route might alter LD homeostasis to alter the outcomes of platinum therapy. Therefore, we utilized a hypothesis-driven (Fig. 6D) web-based survival analyzer with an established algorithm (Chang et al., 2017) to test their correlation. As mentioned previously, LPC is important for LD metabolism; therefore, we examined the hazard ratio (HR) score of clusters of enzymes/genes responsible for the endogenous lipid resources of LDs (that is, the Kennedy pathway) (Fig. 6D, left-hand side; and Expanded Table 2~4) and found very little contribution to overall, taxol-based therapy, and post-surgery EOC patients’ disease progression (3-yr progression-free survival; PFS) (Expanded Tables 2, 3, and 4, respectively). However, there are gatekeeper genes that influence LDLR through LPL and LDLRAP (LDLR associated protein), and consequently through endoplasmic reticulum (ER) AGPS (alkylglycerone phosphate synthase), EPT (CDP-Ethanolamine:DAG ethanolamine phosphotransferase), and PEMT (Phosphatidylethanolamine N-Methyltransferase) for platinum-based therapy PFS in EOC patients (Fig. 6E, and Expanded Tables 2~4). The net HR score of these gatekeeper genes (+269.86) favors poor disease prognosis, which indicates that the LDL/R-route→LD metabolism could reduce platinum therapy efficacy through PC← → LPC conversion. The discrepancies in platinum therapy and other therapeutic regimens in terms of the progression of patients (Expanded Tables 2~4) suggest a unique function of the LDL/R-route in the remodeling of LD lipidome homeostasis in chemotherapy sensitivity. Except for a bioinformatics simulation of the LDL/R-route in LD homeostasis, we measured the platinum concentrations in LDs resulting from various cisplatin treatments. We found that the LPC-liposome-cisplatin treatment could increase the LD-platinum concentration (Fig. 6F). Additionally, the LPC-liposome-cisplatin treatment also facilitates the binding of DNA to form DNA-platinum adducts (Fig. 6G) in MDAH-2774, while the direct treatment of SKOV3 cells with cisplatin was also able to cause the formation of platinum-DNA adducts (Fig. 6H).

**Figure 6.**
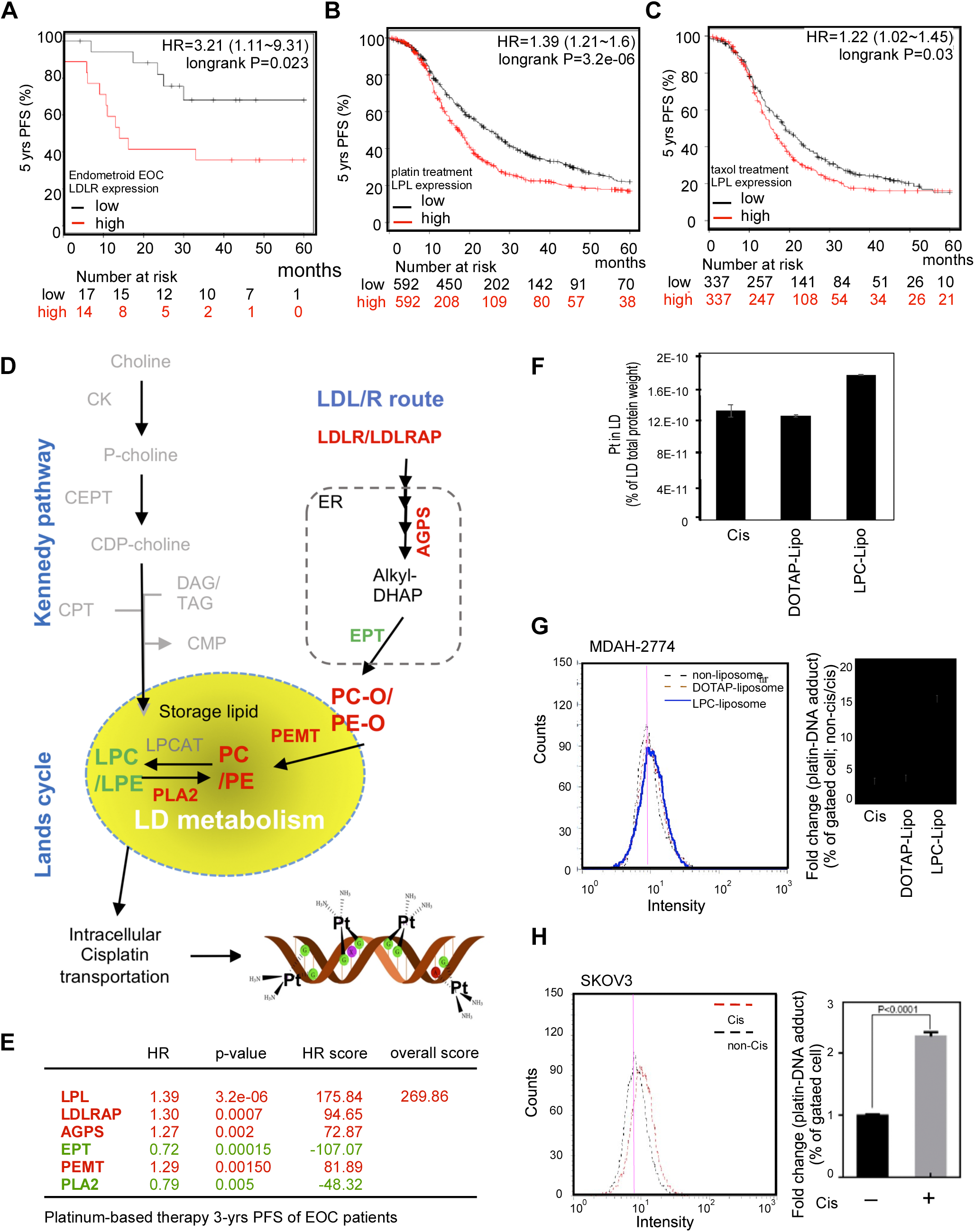
LDL/R-route related to lipid droplet (LD) homeostasis is the biosignature of platinum-based chemotherapy prognosis in EOC patients. **A**. Using the web-based KMplotter survival analyzer to evaluate the impact of LDL/R-route in EOC prognosis, the LDLR expression was found to be positively correlated with the 5-yr progression free survival (PFS) of endometrioid EOC patients. The HR = 3.21, p=0.023. **B** and **C**. The LPL expressions are positively correlated with the 5-yr PFS of EOC patients treated with platinum (**B**; HR=1.39) and taxol (**C**; HR=1.22) therapies; however, the p-value is less for platinum (p=2.3e-06) than for taxol (p=0.03) therapies. **D**. LDL/R-route and Lands cycle for LD metabolism are platinum therapy prognosis biosignatures in EOC patients. Illustration on the left-hand side: Two lipid resources of LD, including the Kennedy pathway and LDL/R-route. Using the web-based KMplotter survival analyzer to evaluate the impact of the Kennedy pathway (grey-colored; Expanded Tables 2~4) showed that it had little influence on the 3-yr PFS of EOC patients. On the other hand, the LDL/R-route to ER→LD metabolism could significantly influence the prognosis of EOC patients receiving platinum therapy. **E**. The table shows the genes with significant impacts. Red-color labeled genes are prognosis promoters, whereas the green-color labeled genes are prognosis suppressors. The sum of the HR score of the overall pathway from the LDL/R-route to the Lands cycle is ~ +2626.05, which indicates a prognosis confounder role of this pathway. **F and G**. The cisplatin encapsulated by LPC-liposome could enhance platinum-DNA binding compared to DOTAP-liposome or cisplatin treatments. The left-hand side: The representative data of flow cytometry measuring platinum-DNA adducts with specific antibodies in MDAH-2774 (**F**; cisplatin insensitive cell) or SKOV3 (**G**; cisplatin sensitive cell) cells. The right-hand side: Quantitation of the flow cytometry results. The Y-axis is the value of fold change (% to gated positive cell when comparing cisplatin/non-cisplatin treated cells). All the data of the in vitro experiments were from at least three independent experiments with consistent results. * indicates a significant difference due to a p-value < 0.05; ** indicates a p-value < 0. 01; and *** indicates a p-value < 0.001.

The prediction of the LDL/R-route by web-based survival analyses has demonstrated the possible mechanisms of LDLR engulfing (to LD), LPC metabolism (via the Lands cycle), and finally the influence of LD homeostasis. In addition, it was shown that the overall activity is related to the prognosis of platinum-based therapy in EOC patients. We further proved this possibility in vitro by testing the effects of LPC-liposome-cisplatin treatment on LD metabolism and the amount of platinum-DNA adduct in EOC cells.

### LPC-liposome as effective cisplatin efficacy booster for cancer therapy

In this study, the LDLR-mediated lipidome reprogramming in EOC cells, e.g., that via LPC, was found to be the platinum-based therapy sensitivity confounder. In order to translate this discovery to a potential therapeutic strategy, we tested whether or not using the LPC-liposome as a cisplatin booster could be generalized to other types of cancer. We first characterized the cisplatin responsiveness in 43 cell lines (carcinoma-derived cancer cells), and then selected 22 cells with high IC 50 values (>100 Expanded Table 5). Because the primary mechanism of action for platinum cytotoxicity occurs via DNA adduction formation, thereby blocking the new synthesis of double-stranded DNA, the sensitive cells should be fast growing ones, in theory. Therefore, the doubling times of the 22 cell lines were characterized in order to produce a shorter list (only those with a doubling time < 50-hrs were selected) of 10 cell lines (Fig. 7A). These 10 cell lines included hepatocellular carcinoma (HCC: HepG2), renal cell carcinoma (RCC: 769-p), pancreatic cancer (PanCa: CFPAC, HPAF-II, AsPC-1, BxPC-3), epithelial ovarian cancer (EOC: MDAH-2774), gastric cancer (GCa: AGS), and cholangiocarcinoma (CC: RBE, SSP25) cell lines, among others. As shown in Fig. 7B, when we tested its effects in MDAH-2774 (endometrioid EOC) and TOV-21G (clear cell EOC) cells, we found that LPC-liposome-cisplatin treatment could efficiently suppress both types of cells with 20 μM of treatment. When we tested 769-p (RCC; clear cell type), HepG2, and AGS (in which a vacuolar morphology is commonly seen) cells, we found that LPC-liposome treatment also dramatically suppresses cell growth (Fig. 7C). Lastly, we tested two difficult to treat types of cancer cells (PanCa and CC), and found that cisplatin in an LPC-liposome-encapsulated form exerted excellent cell growth suppression capacity. This result clearly demonstrated the feasibility of using LPC-liposome as a platinum-based therapy efficacy booster. To sum up, LPC-liposome could potentially be widely used as a cisplatin enhancer in multiple cancer therapies.

**Figure 7.**
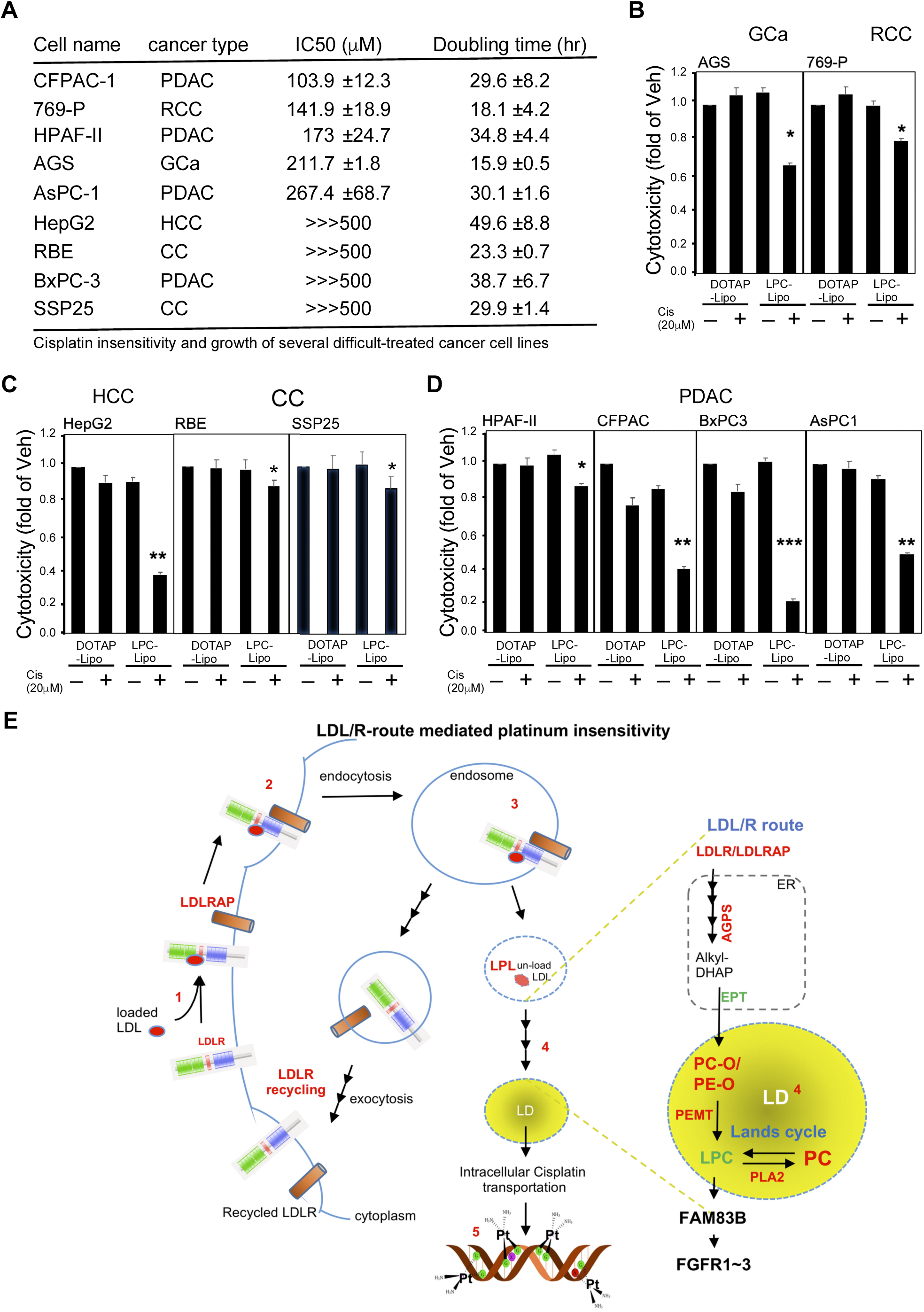
LPC-liposome is an excellent cisplatin efficacy booster in multiple types of cancer cells. **A**. The list of cancer cells with high cisplatin IC 50 values (insensitive cells; IC 50 > 100 μM) that are fast growing (doubling time less than 50 hrs). The types of cancer cells included pancreatic cancer (PanC; CFPAC-1, HPAF-II, AsPc-1, BxPC-3), renal cell carcinoma (RCC; 769-P), epithelial ovarian carcinoma (EOC; MDAH-2774), gastric cancer (GCa; AGS), hepatocellular carcinoma (HCC; HepG2), and cholangiocarcinoma (CC; REB and SSP25) cell lines. **B~D**. The LPC-liposome exhibited excellent cisplatin cytotoxic boosting efficacy in multiple types of cancer, including EOC (**B**), gut cancer (**C**; HCC, GCa, CC), PanC, and RCC (**D**). **E**. Schematic illustration of LDL/R-route-mediated lipidome reprogramming to facilitate LD lipid remodeling and FGFR signaling in platinum-based therapy sensitivity.

## Discussion

In this study, we found that the histological variation of EOC differential expression of the LDL/R-route determines platinum-based therapy insensitivity. The mechanism underlying this could involve the LDLR→LPC →FAM83B →FGFRs regulatory axis, in which lipid metabolism takes place in LDs. The overall findings can be drawn as the illustrative scheme shown in Fig. 7E. Furthermore, the potential impacts of our findings are discussed below:

### Dependency of chemosensitivity and targeting strategies on LDL/R-route-lipidome reprogramming

The effects of LDL/R-route and related lipidome expression on chemosensitivity have previously been hypothesized but had not previously been verified (Huang, Li et al., 2016). With regard to platinum-based therapy, it was previously shown that the latency from initial adjuvant platinum therapy to resistance is around 90~250 days in lung adenocarcinoma (Wu, Si et al., 2015). It has been speculated that the mechanism underlying this resistance could involve cholesterol-induced ABCG2 expression, with the ABCG2 being recognized as a channel for pumping out lipophilic wastes, e.g., platinum chemoagents. Interestingly, LD function is related to lipid transport through ABC proteins (Baldan, Bojanic et al., 2009, Gulati, Balderes et al., 2015). Relatedly, a recent study using transcriptome and metabolomics analyses in NCI-60 cell lines found that lipoprotein uptake is one of the hallmark events in platinum sensitivity (Cavill, Kamburov et al., 2011). As for other chemoagents, other evidence has shown that cholesterol uptake (potentially via LDLR) could be an important event for developing gemcitabine resistance in cases of pancreatic cancer, which has caused such uptake to be considered an excellent therapeutic target (Guillaumond, Bidaut et al., 2015). In this study, we discovered a LDLR→LPC→FAM83B→FGFRs axis for platinum sensitivity. Taking advantage of this discovery, we then implemented the use of LPC-liposome-encapsulated cisplatin as a new chemoagent. The ensuing proof-of-concept experiment detailed in this report demonstrated an excellent cytotoxic capacity of LPC-liposome-cisplatin in multiple cancer types which are insensitive to cisplatin treatments.

Proprotein convertase subtilisin/kexin type 9 (PCSK9) is a soluble member of the mammalian proprotein convertase family of secretory serine endoproteases (Seidah, Benjannet et al., 2003). Circulating PCSK9 is known to degrade LDLR (Lagace, Curtis et al., 2006) via binding on the epidermal growth factor-like repeat A site of LDLR, which causes the recycling activity to be reduced and, thus, causes the degradation of LDLR (Zhang, Lagace et al., 2007). The mechanism-of-action of PCSK-9 inhibitors (such as alirocumab and evolocumab) exerts anti-degrading liver LDLR activity; therefore, these PCSK-9 inhibitors enhance LDLR recycling to reduce circulating LDL levels. In this study, we discovered that the LDL/R-route reduces platinum efficacy. Therefore, one of the strategies for targeting the LDL/R-route consists of reducing systemic LDL levels. Introducing a PCSK-9 inhibitor in the newly developed LPC-liposome-cisplatin for this purpose exhibited great potential. On the other hand, an LDL/R-route-targeting strategy could also include degraded cancer LDLR. One in vitro study showed that PCSK-9 could degrade LDLR by interacting with glypican-3 in liver cancer cells (Ly, Essalmani et al., 2016). The development of PCSK-9 peptide for degrading LDLR (Lagace, 2014) is also a hypothetical strategy for platinum-based therapy.

### Targeting LD lipid remodeling for chemotherapy

Lipid droplets (LDs) are intracellular lipid storage organelles consisting of a core of neutral lipids, e.g., DAG or TAG, and a surrounding monolayer of phospholipids, predominantly phosphatidylcholine (PC) (Moessinger et al., 2014). The accumulation of LDs is a well-recognized hallmark of cancer; however, the role of LDs in cancer is still unknown. The metabolism of LDs relies on the conversion of PC to LPC (via the Lands cycle), which is dynamically balanced by LPCAT1/2/3 (lysophosphatidylcholine acyltransferase 1/2/3) and PLAs (phospholipase A) (Chen, Kazachkov et al., 2007, Moessinger et al., 2014, Moessinger, Kuerschner et al., 2011). These lipid resources can be both endogenous and exogenous with endogenous lipids potentially contributed via the Kennedy pathway (Gibellini & Smith, 2010). In the bioinformatics analyses and in vitro validation conducted in this study, we observed significant impacts of the LDL/R→sER→LD route, but not the Kennedy pathway, on the prognosis of EOC patients receiving platinum-based therapy. This study revealed, for the first time, the importance of exogenous lipids for reprogramming the cellular lipid composition for LD metabolism. There are also some other recent studies that support this claim. For example, one report studying a colorectal cancer mouse model found that LPCAT2 contributed to LD accumulation, resulting in chemotherapy resistance (Cotte, Aires et al., 2018). Aside from chemosensitivity, a 2018 study by Wang et al. 2018 reported that LPCAT3 could cause phospholipid remodeling and promote stem cell proliferation, which is related to colorectal cancer tumorigenesis (Wang, Rong et al., 2018).

Meanwhile, although a few previous reports have discussed the roles of LDs in cancers, none have discussed the potential of targeting LD as a cancer therapy. In this study, the experimental treatment of LPC with cisplatin did not alter insensitive cells, but LPC-liposome-cisplatin treatment exhibited excellent growth suppressing efficacy. The possible mechanism consist of the LPC-liposome (a mixture of LPC and cholesterol) mimicking the structure of LDs and then fusing with LDs to remodel their lipid composition. There is great interest in exploring this possibility further for the purposes of developing new drug delivery methods. Furthermore, this result suggests the great potential of targeting LDs for cancer therapy. Relatedly, the concept of nano-droplet "adiposome" (Wang, Zhou et al., 2016) has been proposed as a tool for drug delivery, although this concept requires future validation.

In conclusion, this study discovered that LDLR expression is a platinum chemosensitivity confounder. Furthermore, the novel LDLR→LPC→FAM83B→FGFRs regulatory axis revealed by the trans-omics analyses conducted in this investigation might explain platinum chemosensitivity discrepancies. Finally, LDLR-altered LD homeostasis was found to contribute to platinum sensitivity, which suggests the potential value of targeting LDs with LPC-liposome-cisplatin treatment.

## Materials and Methods

### Patient cohort

The paraformaldehyde-embedded EOC tissue samples analyzed in this study were obtained from patients diagnosed with EOC from 2008 to 2013 at China Medical University (Taichung, Taiwan). The patients were identified from a single cohort registered in the Cancer Registry Database of the hospital, and the gynecological pathology of each patient was classified according to the World Health Organization pathology classification. Access to the tissue samples was approved by the Internal Review Board of China Medical University Hospital (# DMR101-IRB2-276; and CMU105-REC3-122(CR1)). The EOC subtypes selected from patient chart overview, and then confirmed based on two boarded pathologists diagnosed on H&E stained paraffin sectioned slides to exclude ambiguity or mix histology between subtypes.

### Immunohistochemistry and quantitation of staining score

In general, the histological studies were performed as described in previous studies (Chen, Bao et al., 2018, Hung, Chang et al., 2014) with some modifications. For general histologic inspection, we treated the tissue sections (2 μM) with hematoxylin and eosin or stained the sections with antibodies specific for LDLR and LPL with an ABC kit (Vector Laboratories) to enhance the staining signals. Staining intensity was scored according to the Allred scoring system (Hammond, Hayes et al., 2010, Nose, Sugio et al., 2009) and our previous work (Lai, Yeh et al., 2016). The proportion of cells that stained positive for LDLR and LPL was graded using a five-point scale according to the proportion of positive cancer cells (1: < 1/100; 2: 1/100 to 1/10; 3: 1/10 to 1/3; 4: 1/3 to 2/3; and 5: > 2/3). The intensity of staining was also graded on a five-point scale as follows: 1: none; 2: weak; 3: intermediate; 4: mid-strong; 5: strong. The proportions and intensity scores were next sum, averaged, and then compared with the associated histological reports. The slides were independently examined by three coauthors (WC Chang, MD; HS Wang; and YP Ho) who were blinded to the clinicopathological data. When there was a discrepancy (i.e., a score difference > 3) between the scores given by the slide reviewers, the pathologists reassessed the slides using a double-headed microscope, and a consensus was reached. Finally, associations between the scores and the clinical data were investigated by another coauthor (YC Hung, MD).

### Reagents, cell culture, and lentiviral-based gene delivery

Cells were maintained in various culture media (depending on the culture requirements) with 10% FCS (Invitrogen), 1% L-glutamine, and 1% penicillin/streptomycin as described previously (Chang et al., 2017). The HEK293T (ATCC; HTB52) and EOC (MDAH-2774; SKOV3, HTB-77; OVCAR3, HTB-161; ES2, CRL-1978; TOV-112D, CRL-11731; TPOV-21G, CRL-11730) cell lines were purchased from ATCC; the head-neck squamous cell carcinoma (HNSCC: OECM1, FaDu, SAS) cell lines were the courtesy of Professor Kou-Juey Wu of CMU; the PanCa (CFPAC-1, HPAF-II, ASPC-1, BxPC-3) cell lines were the courtesy of Professor Wen-Hwa Lee of CMU; the HCC (Tong, HCC36, Huh7, HepG2) cell lines were kindly provided by Dr. YS Jou of the Academia Sinica; the GCa (AGS, MKU-1, SC-M1) cell lines were purchased from the Food Industry Research and Development Institute in Taiwan (BCRC purchase number: 60210); the RCC (769-p) cell line was kindly provided by Professor Chawnshang Chang of the University of Rochester, NY, USA; and the CC (H1, RBE, SSP25) cell lines provided by Chiung-Kwei Huang of Brown University.

The following antibodies were used: LDLR (for IHC: GeneTex, GTX61553; for immunoblot: Santa Cruz, sc-373830), LPL (Santa Cruz, sc-32885), VEGFR (Santa Cruz, sc-6251), phospho-FAK (Cell Signaling Technology, #3283), FAK (Cell Signaling Technology, #3285), MEK1/2 (Cell Signaling Technology, #8727), Actin (Santa Cruz, sc-47778), and tubulin (Abcam, ab-6046). The following chemicals were also used: cisplatin (Sigma-Aldrich, P4394), DOTAP (1,2-dioleoyl-3-trimethylammonium-propane; Avanti, 890890P), and lyso-phosphatidylcholine (Avanti, 855675P),

### Lentiviral-based gene delivery

(Ma, Hsu et al., 2012, Ma, Jeng et al., 2014): LDLR knockdown or overexpression clones were engineered by the stable transfection of human LDLR cDNA (pLenti-C-mGFP-LDLR, RC200006L2; OriGENE, Rockville, MD, USA) or pLKO.1-shLDLR (targeting sequence shown in Expanded Table 6) and then selected after exposure to puromycin (10 μM) for a period of time. The pLKO-shLuciferase, shLDLR, shFAM83B (Expanded Table 6) plasmids were obtained from the National RNAi Core Facility Platform (Institute of Molecular Biology/Genome Research Center, Academia Sinica, supported by the National Core Facility Program for Biotechnology; grant NSC107-2319-B-001-002). The pBabe and pWPI (Addgene) vector-based AR cDNA expression plasmids were constructed as previously reported (Ma, Hsu et al., 2008). The lentiviral production and infection procedures used in this study followed those reported previously (Ma et al., 2012). In brief, psPAX2 (packaging plasmid) and pMD2G (envelope plasmid) (Addgene) were co-transfected into HEK293T cells. We then harvested virus-containing media to infect the HCC cells. The GFP+ cell populations, as determined by flow cytometry analysis (BD LSR II Flow Cytometry), were used to test the infection efficiencies.

### Colony formation, cytotoxic measurement, and IC 50 values

Colony-forming assays were performed as previously reported (Chen, Chang et al., 2014). Briefly, 1×10^4^ cells/dish were seeded onto 3.5-cm plates with DMEM in 10% FBS with various treatments for 7 days. After the treatments, 1/3 of the total volume of the 10% formaldehyde solution was added to fix the cells, which were then allowed to stain with Crystal Violet for 5 mins. After being washed with PBA, the colonies were photographed.

For the cell viability assay, cells were seeded in 96-well plates (5 × 10^3^ cells/well) and incubated overnight for attachment, and were then treated with indicated doses of drugs in normal media for 48 hr. After the treatments, the media were replaced with MTT (0.5 mg/ml) at 37°C for at least 1 hr. After the removal of excess WST-1 (Sigma-Aldrich), the colorimetric absorbance of the cells at 490 nm was read. The readings of the measured values of 50% inhibition concentration (IC 50) (Chou, 2010) for each drug were determined by CalcuSyn software (Chou & Talalay, 1984) (BioSoft).

### Experimental animal and xenograft implantation tumor model

Athymic nude female mice aged 6–8 weeks old *(Foxn1^nu^)* were purchased from the NLAC (National Laboratory Animal Center), Taiwan. Subcutaneous implantation of 1 × 10^6^ cells/100 μl PBS and matrigel (1:1) in both flanks was performed on each mouse. The mice were then randomly divided into experimental groups as the tumors grew to 500-mm^3^, and the size of each tumor was measured twice/week. The mice were treated with/without cisplatin (6 mg/kg/mice; or equal volume of phosphate buffered saline; I.P., every other day for 4 wks). The mice were then sacrificed and the tumors were harvested. All the animal studies were performed under the supervision and guidelines of the China Medical University Animal Care and Use Committee (#CMOIACUC-2018-089).

### RealTime RT-PCR and primers

The protocol for detecting mRNA expression followed that detailed in a previous publication (Chiang, Wang et al., 2017) with some modifications. Total RNA was isolated from the tissue using the Trizol™ reagent (Invitrogen, USA) according to the manufacturer’s protocol. The mRNA levels of various genes were measured by qPCR using the Bio-Rad CFX 96 sequence detection instrument. The levels of mRNA were normalized with GAPDH mRNA. The SYBR probe (Bio-Rad, USA) was used as the fluorogenic probe to determine the threshold cycle (Ct), and the forward and reverse primers are shown in Expanded Table 6.

### Lipid profiling for lipidome analysis

After the cells (1500 cells/μl × 300μl) were washed with Ca^2+^/Mg^2+^-free PBS, the lysates were then subjected to lipid profiling executed by Lipotype GmbH (Dresden, Germany) (Ejsing, Sampaio et al., 2009, Levental, Surma et al., 2017, Sampaio, Gerl et al., 2011). Lipidomes were prepared from at least three replicates of each sample for all the experiments using the following procedures.

Nomenclature. The following lipid names and abbreviations were used: Cer, ceramide; Chol, cholesterol; DAG, diacylglycerol; HexCer, glucosyl/galactosyl ceramide; PA, phosphatidic acid; PC, phosphatidylcholine; PE, phosphatidylethanolamine; PG, phosphatidylglycerol; PI, phosphatidylinositol; PS, phosphatidylserine; and their respective lysospecies lyso-PA (LPA), lyso-PC (LPC), lyso-PE (LPE), lyso-PI (LPI), and lyso-PS (LPS); and their ether derivatives PC O-, PE O-, LPC O-, and LPE O-; SE, sterol ester; SM, sphingomyelin; SLs, sphingolipids; and TAG, triacylglycerol. Lipid species were annotated according to their molecular composition as follows: [lipid class]-[sum of carbon atoms in the fatty acids]:[sum of double bonds in the fatty acids];[sum of hydroxyl groups in the long-chain base and the fatty acid moiety] (for example, SM-32:2;1). Where available, individual fatty acid compositions following the same rules were given in brackets (for example, 18:1;0–24:2;0).

Lipid extraction for MS lipidomics. Lipids were extracted using a two-step chloroform/methanol procedure. Samples were spiked with internal lipid standard mixture containing cardiolipin (CL), 16:1/15:0/15:0/15:0; Cer, 18:1;2/17:0; DAG, 17:0/17:0; HexCer, 18:1;2/12:0; LPA, 17:0; LPC, 12:0; LPE, 17:1; LPG, 17:1; LPI, 17:1; LPS, 17:1; PA, 17:0/17:0; PC, 17:0/17:0; PE, 17:0/17:0; PG, 17:0/17:0; PI, 16:0/16:0; PS, 17:0/17:0; cholesterol ester (CE), 20:0; SM, 18:1;2/12:0;0; TAG, 17:0/17:0/17:0; and Chol. After extraction, the organic phase was transferred to an infusion plate and dried in a speed vacuum concentrator. Each first-step dry extract was resuspended in 7.5 mM ammonium acetate in chloroform/methanol/propanol (1:2:4, v/v/v), and each second-step dry extract was resuspended in a 33% ethanol solution of methylamine/chloroform/methanol (0.003:5:1, v/v/v). All liquid handling steps were performed using the Hamilton Robotics STARlet robotic platform with the Anti-Droplet Control feature for organic solvent pipetting.

MS data acquisition. Samples were analyzed by direct infusion on a Q-Exactive mass spectrometer (Thermo Scientific) equipped with a TriVersa NanoMate ion source (Advion Biosciences). Samples were analyzed in both the positive and negative ion modes with a resolution of 280,000 at m/z = 200 for MS and 17,500 for MS/MS experiments in a single acquisition. MS/MS was triggered by an inclusion list encompassing corresponding MS mass ranges scanned in 1-Da increments. Both MS and MS/MS data were combined to monitor CE, DAG, and TAG ions as ammonium adducts; PC and PC O- as acetate adducts; and CL, PA, PE, PE O-, PG, PI, and PS as deprotonated anions. Only MS was used to monitor LPA, LPE, LPE O-, LPI, and LPS as deprotonated anions; Cer, HexCer, SM, LPC, and LPC O- as acetate adducts; and Chol as an ammonium adduct of an acetylated derivative (Surma, Herzog et al., 2015).

Data analysis and post-processing (Klose, Surma et al., 2013). Lipid identification using LipotypeXplorer (2) was performed on unprocessed mass spectra. For the MS-only mode, lipid identification was based on the molecular masses of the intact molecules. The MS/MS mode included the collision-induced fragmentation of lipid molecules, and lipid identification was based on both the intact masses and the masses of the fragments. Prior to normalization and further statistical analysis, the lipid identifications were filtered according to mass accuracy, occupation threshold, noise, and background. Lists of identified lipids and their intensities were stored in a database optimized for the particular structure inherent to lipidomic datasets. The intensity of lipid class-specific internal standards was used for lipid quantification (Liebisch, Binder et al., 2006). The identified lipid molecules were quantified by normalization to a lipid class-specific internal standard. The amounts in p-moles of individual lipid molecules (species of subspecies) of a given lipid class were summed to yield the total amount of the lipid class. The amounts of the lipid classes may be normalized to the total lipid amount, yielding mol. % per total lipids.

### Lipidomic data processing

The lipid profiling data for each sample was scale normalized by the total amount of lipid. Those lipids with at least a two-fold change between the LDLR knockdown and control cells were identified as the lipids that were significantly regulated by LDLR. Then, Fisher’s exact test was conducted to test the enrichment of the significantly regulated lipids in each lipid class (such as PC, PE, and LPC).

### TCGA-database DriverDB.v2 and KMplotter meta-analysis for cancer survival analysis, and algorithm for hazard ratio scoring

The previously developed DriverDB (Cheng, Chung et al., 2014, Chung, Chen et al., 2015), a database that incorporates more than 9500 cancer-related RNA-seq datasets and more than 7000 exome-seq. datasets from TCGA, was used in this study. In the DriverDB, there are 420 primary tumors and 37 adjacent normal tissues (including 34 normal-tumor pairs) in the EOC dataset from TCGA. For the survival analysis of TCGA data, KaplanMeier survival curves were drawn and the log-rank test was performed to assess the differences between the patient groups stratified by the median of gene expression. A p-value of less than 0.05 was considered statistically significant.

With regard to the web-based KMplotter platform used for the evaluation of the hazard ratio (HR) scores of the pathways (cluster of genes) with respect to patient survival, the following previously established formula was used (Chang et al., 2017, Chang, Huang et al., 2016):

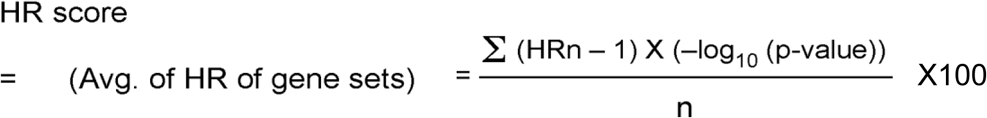

In order to evaluate the impact of each gene, the absolute value of the HR for that gene minus 1 was calculated. To adjust for the effects of the genes, the HR value for each gene was multiplied by negative log_10_(p-value) to balance the importance of the genes. The summed score was then divided by the number of genes and multiplied by 100 to obtain the HR score, or the average HR of all the genes.

### Preparation and characterization of LPC-liposome

The liposome was prepared by thin-layer hydration (Jang, Kim et al., 2013) followed by application of the membrane protrusion method (Ong, Chitneni et al., 2016) with some modifications. First, we hydrated a mixture of DOTAP (mw=698.5 g) or LPC (mw=495.63 g) : cholesterol (mw=386.6 g)= 1:1 (molar ratio) with purified water (Milli-Q Plus, Millipore, Bedford, MA, USA). The mixture was then incubated at 65°C for an hour, and then sonicated in a 65°C water-bath for 30 min. The mixture was then subjected to membrane protrusion (mini-Extruder, Avanti Polar Lipid, Ltd.) with a 200-nm pore size membrane (Avanti Polar Lipid, Ltd.), being extruded 20 times to form pre-liposome. The pre-liposome was then subjected to protrusion with a 100-nm pore size membrane, being extruded another 15 times. The liposome size and size distribution were then determined by photon correlation spectroscopy (Zetasizer Nano-ZS90, Malvern Instruments Ltd., Malvern, Worcestershire, UK). The 10-μL liposomes were dispersed with 500 μL purified water in a low volume disposable sizing cuvette. The particle size and size distribution were measured in terms of ZAve and polydispersity index (PDI), respectively. The LPC-liposome-encapsulated cisplatin size and distribution are shown in EV.3.

### RNA sequencing for transcriptome analysis

RNA-Seq libraries were prepared using the Agilent SureSelect RNA Library Kit and were sequenced using Illumina Hiseq4000 150-nt PairedEnd to produce the reads (~25 million reads per sample). The reads were aligned with TopHat 2.0.13 to GRCh38 with default parameters and then were assembled by Cufflink 2.2.1 using Ensembl v79 annotations. Gene expression was measured in fragments per kb of exon per million fragments mapped (FPKM). For the differentially expressed genes regulated with LDLR, we performed functional enrichment analysis, as described in our previous studies (Cheng et al., 2014, Chung et al., 2015), to interpret their biological functions.

### Cisplatin-DNA adduct measurement with flow cytometry

Platinum-DNA adduct measurement was described previously. (Lundholm, Haag et al., 2013) In brief, cells were subjected to various treatments for 24 hrs to form cisplatin-DNA adducts. The cells were then harvested with PBS containing 0.1% Triton X-100 and fixed with 70% ethanol. Next, the cells were washed with PBS and stained with anti-cisplatin modified DNA antibody [CP9/19] (Abcam; 1:1000 dilution) overnight at 4°C. The cells were then stained with goat anti-rat FITC-conjugated secondary antibody for 2 hr. The signals of the cisplatin-DNA adducts were then detected by flow cytometry (BD Biosciences). The data were analyzed using the FCS Express v3.0 software (De Novo Software).

### Quantitation of the uptake of cisplatin in cells

The method of platinum quantitation in cells was modified from a previous study. (Corte-Rodriguez, Espina et al., 2015) The platinum levels were detected and measured by ICP-MS analysis. Cells were treated with 65% nitric acid (0.5 ml) and 30% H2O2 (0.5 ml) for digestion, followed by heating up to 80°C for one hour. After digestion, 25% ammonia water (0.5 ml) was added into the samples to neutralize the excess nitric acid. The samples were then diluted with up to 4 ml of ultrapure water. The measurement of platinum levels was conducted using an ICP-MS Agilent 7900 system (Agilent Technologies) in a certified laboratory (Super Micro Mass Research and Technology Center, Cheng Shiu University)

#### Statistics

The Student *t*-test or the chi-square analysis was used to identify significant differences between groups or categorical variables. A p-value less than 0.05 was considered significant. All data are reported as the mean ± standard error of the mean (SEM).

## Acknowledgements

We appreciated Dr. Keen Chung for instructing the liposome preparation for drug delivery. We also appreciated technical support by Lipotype GmbH (Dresden, Germany) and Taiwan-BioActive Lipid Ltd. Co. (Taiwan) on lipidome profiling and bioinformatics analysis. Thanks also to Professor Chawnshang Chang of the University of Rochester and Professor Chiung-Kwei Huang of Brown University for generosity to provide and verify RCC and cholangicarcinoma cell lines.

## Authors’ contributions

WC Chang collected the clinical samples, analyzed the data, and drafted the manuscript. HC Wang performed the animal experiment and drafted the methodology section, while WC Cheng performed the bioinformatics analysis. JC Yang performed HPLC for platinum measurement and assisted with the in vitro experiment. WM Chung and YP Ho helped with the immunohistochemistry study. YC Hung supported the study, collected the clinical sample, and edited the manuscript. W-L Ma developed the concept, coordinated the research project, support the project, and edited/approved the final version of the manuscript.

Availability of data and materials: All datasets are available from the corresponding author upon reasonable request.

## Conflict of interest

All the authors claimed no interests conflict.

## Notes

**Grant funding**: This study was supported in part by grants from the Taiwan Ministry of Science and Technology (MOST104-2628-B-039-001-MY4; MOST106-2221-E-039-011-MY3); National Health Research Institute (NHRI-EX107-10705BI). This study was also supported in part by China Medical University/Hospital (CMU107-S-05; CMU107-TC-02; DMR-108-080; DMR-CELL-17014; DMR-CELL-1806; DMR-108-079).

Conflict of interest statement: All the authors in this work claimed no interests conflict.

